# Perfect adaptation of CD8^+^ T cell responses to constant antigen input over a wide range of affinity is overcome by costimulation

**DOI:** 10.1101/535385

**Authors:** Nicola Trendel, Philipp Kruger, Stephanie Gaglione, John Nguyen, Johannes Pettmann, Eduardo D Sontag, Omer Dushek

**Author notes:** Corresponding author. Omer Dushek, Sir William Dunn School of Pathology, University of Oxford, South Parks Road, Oxford, OX1 3RE, United Kingdom, E. Equal contribution (Nicola Trendel, Philipp Kruger, Stephanie Gaglione).

## Abstract

Maintaining and limiting T cell responses to constant antigen stimulation is critical to control pathogens and maintain self-tolerance, respectively. Antigen recognition by T cell receptors (TCRs) induces signalling that activates T cells to produce cytokines and also leads to the downregulation of surface TCRs. In other systems, receptor downregulation can induce perfect adaptation to constant stimulation by a mechanism known as state-dependent inactivation that requires complete downregulation of the receptor or the ligand. However, this is not the case for the TCR, and therefore, precisely how TCR downregulation maintains or limits T cell responses is controversial. Here, we observed that *in vitro* expanded primary human T cells exhibit perfect adaptation in cytokine production to constant antigen stimulation across a 100,000-fold variation in affinity with partial TCR downregulation. By directly fitting a mechanistic model to the data, we show that TCR downregulation produces imperfect adaptation, but when coupled to a switch produces perfect adaptation in cytokine production. A prediction of the model is that pMHC-induced TCR signalling continues after adaptation and this is confirmed by showing that, while costimulation cannot prevent adaptation, CD28 and 4-1BB signalling reactivated adapted T cells to produce cytokines in a pMHC-dependent manner. We show that adaptation also applied to 1st generation chimeric antigen receptor (CAR)-T cells but is partially avoided in 2nd generation CARs. These findings highlight that even partial TCR downregulation can limit T cell responses by producing perfect adaptation rendering T cells dependent on costimulation for sustained responses.

## Introduction

T cell activation is critical to initiate and maintain adaptive immunity. It proceeds by the recognition of peptide major-histocompatibility complex (pMHC) antigens by T cells using their T cell receptors (TCRs). TCR/pMHC binding induces signalling pathways that can activate T cells to directly kill cancerous or infected cells and to se-crete a range of cytokines (1). When T cells are confronted with persistent or constant pMHC antigens, maintaining responses to foreign or altered-self pMHC (in chronic infections and cancers (2)) can be just as important as lim-iting responses to self pMHC (e.g. adaptive tolerance (3)). Like other surface receptors, the TCR is downregulated from the surface of T cells upon recognition of pMHC ligands (4). Precisely how TCR downregulation controls T cell responses to constant pMHC antigen stimulation remains controversial.

In other cellular systems, receptor downregulation can induce biological adaptation to constant ligand stimulation (5). Adaptation is defined by the ability of a system to display transient responses that return to baseline when presented with constant input stimulation. The process is known as perfect (or near-perfect) when the baselines before and after stimulation are similar and is imperfect otherwise. Systematic network searches have identified two key mechanisms of adaptation; negative feedback loops (NFLs) and incoherent feedforward loops (IFFLs) (6, 7). At a molecular level, these mechanisms are implemented by surface receptors, signalling pathways, and transcriptional networks (5, 8, 9). In the case of receptor tyrosine kinases (RTKs), G-protein coupled receptors (GPCRs), and ion channels, the common underlying mechanism is effectively an incoherent feedforward termed state-dependent inactivation (5, 7, 9). This mechanism relies on receptors becoming inactivated (i.e. no longer able to signal) after sensing the ligand by, for example, receptor downregulation. Perfect adaptation can be observed when all receptors are downregulated (Fig. 1A) or if all the ligand is removed by the downregulation of receptor/ligand complexes (Fig. 1B).

**Figure 1:**
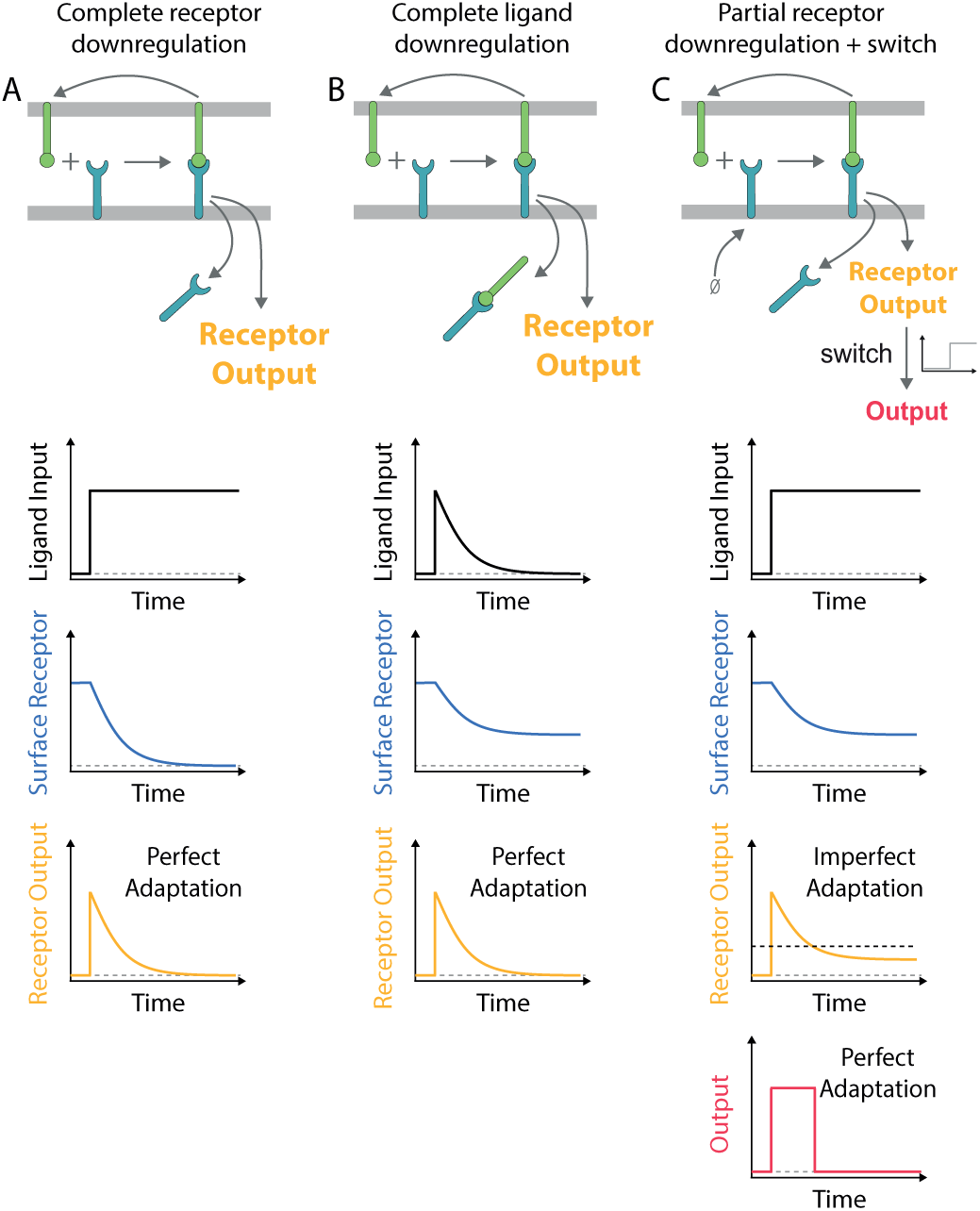
Mechanisms of perfect adaptation based on receptor downregulation. A) Perfect adaptation can be observed if the ligand induces the downregulation of all receptors. This mechanism requires that the re-expression of the receptor on the surface is negligible on the timescale of adaptation. B) Alternatively, perfect adaptation can also be observed with partial receptor downregulation if all ligand is removed by the downregulation of receptor-ligand complexes. This mechanism requires that the receptors are in excess of the ligand (shown) or that receptors are re-expressed on the adaptation timescale (not shown) so that all ligand is removed. C) Receptor output exhibits imperfect adaptation in the model in panel A if the receptor is replenished at the cell surface. In this case, perfect adaptation can be observed if a switch is introduced downstream of the receptor (threshold indicated by dashed horizontal line). In all schematics, the ligand input represents the concentration of ligand available to bind receptor (not including internalised ligand). The mechanism of adaptation by receptor downregulation is a subset of the more general mechanism of state-dependent inactivation (5, 9), which is effectively an incoherent feedforward (7).

The conditions for perfect adaptation exhibited by other receptors are not readily applicable to the TCR. First, the complete downregulation of the TCR is not commonly observed nor is it required for T cell activation (10–14). Second, the complete removal of the pMHC ligand has not been reported although there are reports that some pMHC can be internalised by T cells (15). Instead, individual pMHC ligands have been shown to serially engage and downregulate many TCRs (16) and, on the timescale of hours, they can sustain TCR signalling to induce digital cytokine production (17).

Although TCR downregulation does not appear to meet the criteria for perfect adaptation, it has been suggested to play an important physiological role in limiting T cell responses (18–22). This concept is supported by studies showing that defects in TCR downregulation lead to hyper-responsive T cells with a loss of tolerance to persis-tent self-antigens resulting in autoimmune phenotypes (23–27) and this is associated with sustained early TCR signalling (28, 29). However, studies where transgenic mice were challenged with peripheral antigens came to inconsistent conclusions, with some investigators reporting near-complete TCR downregulation as the mechanism of tolerance (20, 30–33), while others concluded that TCR downregulation did not play a role in tolerance because overt downregulation was not observed (34–38). Therefore, it would seem that complete TCR downregulation is not necessary for adaptation tolerance (3).

Maintaining T cell responses is critical in adoptive therapies where T cells, produced by *in vitro* expansion, are transferred into cancer patients and migrate into tumour microenvironments with chronic cancer antigens (39). These T cells are often armed with affinity-enhanced TCRs or synthetic chimeric antigen receptors (CARs) that re-direct them to tumour cell antigens. Similarly to TCRs, CARs are downregulated as a function of antigen con-centration and initially lower levels render T cells less responsive (40–44). How CAR and TCR downregulation shapes the response of these clinically relevant T cells is poorly understood.

Here, we investigated how constant antigen stimulation regulates responses of clinically relevant *in vitro* expanded primary human CD8^+^ T cells. We observed perfect adaptation in cytokine production, whereby production stops after an initial release, over a 100,000-fold variation in antigen affinity with partial TCR downregulation. Mathe-matical modelling shows that TCR downregulation produces imperfect adaptation, but when coupled to a switch, lead to perfect adaptation in cytokine production. A model prediction is that TCR downregulation reduces, but does not abolish, TCR signalling below the threshold for sustained cytokine production. Consistent with this prediction, we show that adapted T cells are reactivated to produce cytokines by ligation of the costimulatory receptors CD28 and 4-1BB and importantly, this reactivation remained pMHC-dependent. Lastly, we show that adaptation is partially avoided in CAR-T cells that incorporate costimulation within the synthetic receptor. Therefore, adaptation can severely limit the production of cytokines in adoptive transfer therapies and have important implications for the design of adaptation-resistant TCR and CAR constructs.

## Results

### Perfect adaptation of T cell responses to constant antigen over large variation in concentration and affinity

Using a standard adoptive therapy protocol (45), we generated *in vitro* expanded primary human CD8^+^ T cells ex-pressing the therapeutic affinity-enhanced c58c61 TCR (46), which recognises the NY-ESO-1_157-165_ cancer testes antigen peptide bound to HLA-A2 (Fig. 2A, Materials & Methods). In order to allow for constant antigen presentation and to isolate its effects, T cells were stimulated by recombinant pMHC ligands on plates (47–49). This system has been widely used to isolate the effects of specific ligands and to precisely control the duration of stimulation (50).

**Figure 2:**
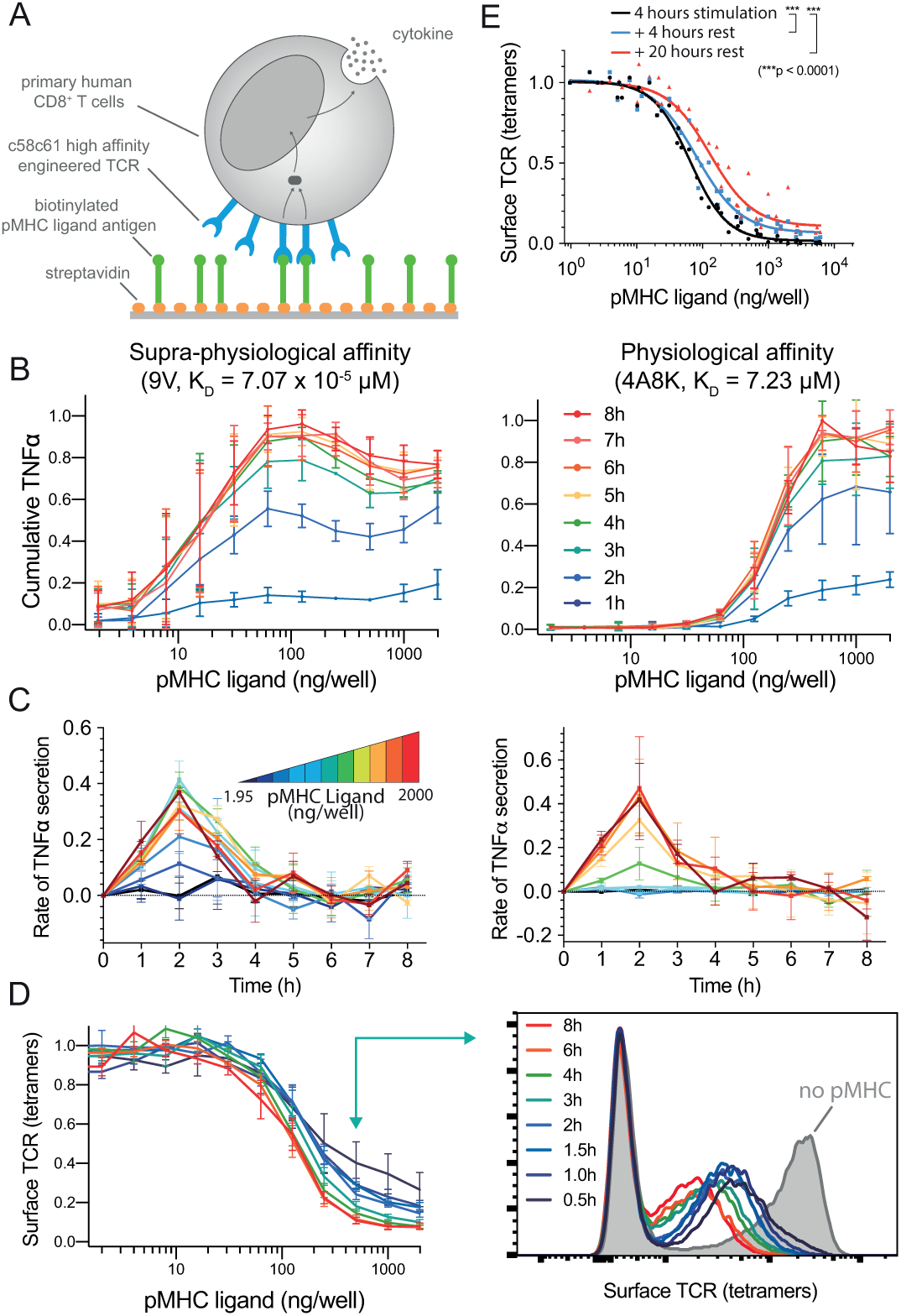
Perfect adaptation of T cells to constant pMHC ligand stimulation over large variation in affinity and concentration, and with proportional downregulation of the TCR. A) Primary human CD8^+^ T cells expressing the c58c61 TCR were stimulated using recombinant pMHC immobilised on plates with supernatant cytokine and surface TCR levels measured (see Materials & Methods). B) Cumulative TNF-α over the concentration of 9V (left) or 4A8K (right) pMHC for 1-8 hours. Mean and SD of 3 independent repeats. C) Data in panel B expressed as a rate of TNF-α secretion over time. D) Surface TCR expression measured using pMHC tetramers in flow cytometry for 4A8K (left) with a representative histogram (right). Mean and SD of 3 independent repeats. E) Recovery of surface TCR was measured by stimulating T cells for 4 hours to induce downregulation (black line) followed by transfer to empty plates without pMHC for 4 (blue) or 20 (red) hours before measuring surface TCR levels. The supernatant levels of MIP-1β, IFN-γ, and IL-2 along with raw data prior to averaging is summarised in Fig. S1-2 and single-cell intracellular cytokine staining in Fig. S3.

T cells stimulated by the high-affinity pMHC antigen (9V, K_D_= 70.7 pM) exhibited perfect adaptation with the secretion of TNF-α stopping after 3 hours (Fig. 2B-C, left column). This adaptation was observed at all antigen concentrations tested. Within this range, high concentrations induced an earlier decline in the rate of TNF-α secretion starting at 2 hours. This resulted in a bell-shaped dose-response curve, which has been previously observed in this system (47) and in other experimental systems (51).

Given that this adaptation was observed with a supra-physiological antigen affinity, we could not exclude the possibility that it was an uncharacteristic response to an excessive antigen signal. We therefore repeated the experiment with a physiological affinity pMHC (4A8K, K_D_= 7.23 µM). Although a higher concentration was required to initiate TNF-α production, the adaptation phenotype was kinetically identical (Fig. 2B-C, right column). We also observed the adaptation phenotype for IL-2, MIP-1β, and IFN-γ (Fig. S1-2), with IFN-γ adapting on a longer timescale as confirmed by transfer experiments (see below; Fig. S8). This distinct behaviour for IFN-γ could result from a subset of T cells being pre-programmed to produce IFN-γ after initially producing TNF-α (52).

The constant level of supernatant cytokine may be established by a balance of uptake with continued secretion or by a stop in secretion. Using single-cell intracellular cytokine staining, we observed that fewer T cells stained positive for TNF-α beyond 3 hours (Fig. S3) suggesting that production stops, consistent with a recent report (53). Moreover, replacing the media and transferring T cells to new plates does not induce cytokine production (see below; Fig. 4C-D, transfer to pMHC & Fig. S7,S8). Collectively, this shows that cytokine production stops in response to constant pMHC ligand stimulation.

**Figure 4:**
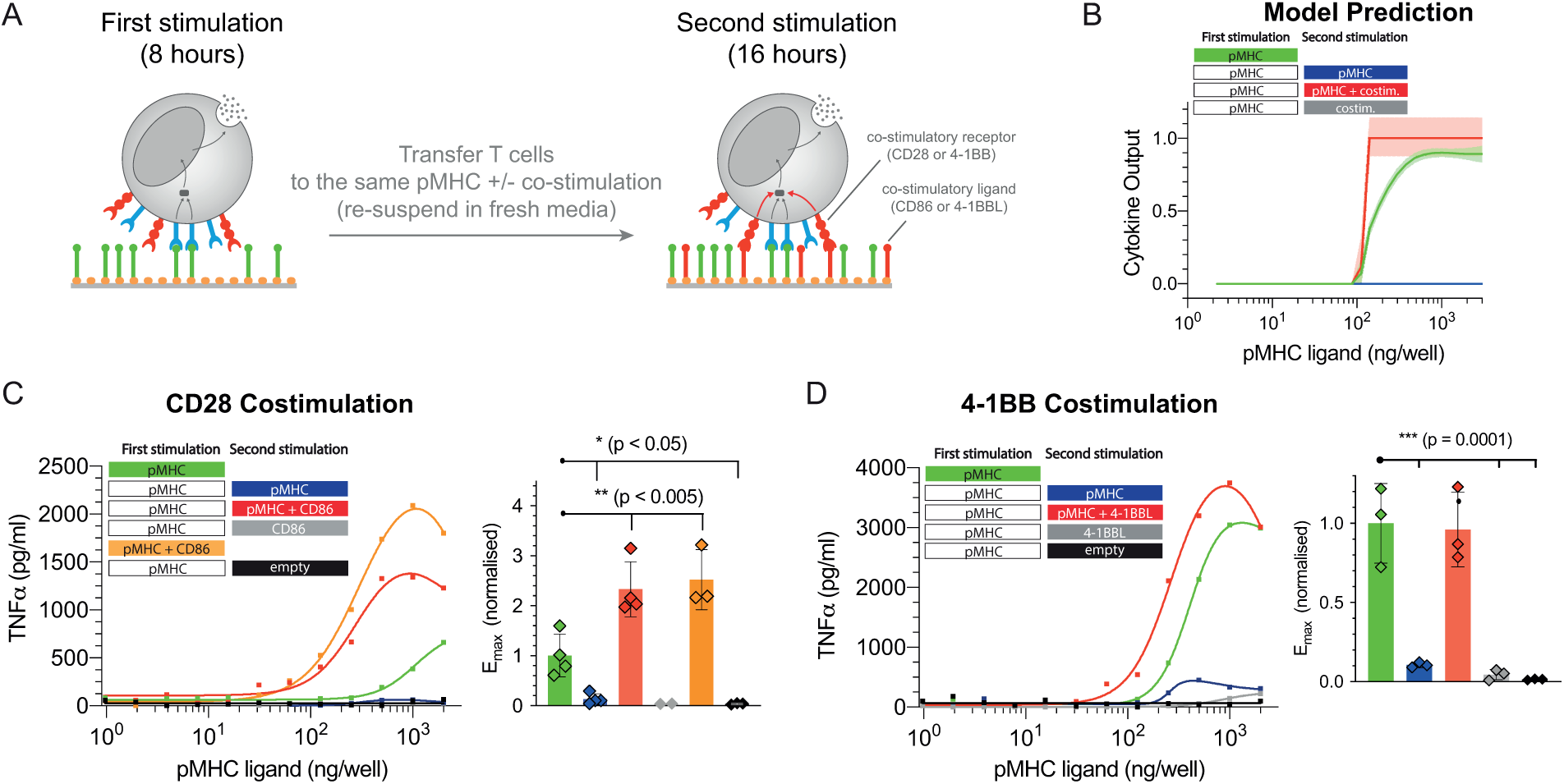
Adapted T cells can be reactivated by CD28 or 4-1BB costimulation. A) Schematic of the experiment showing that T cells were first stimulated for 8 hours before being transferred for a second stimulation for 16 hours with either antigen alone, costimulation alone, or antigen and costimulation. B) Predicted cytokine production by the mathematical model where costimulation is assumed to lower the threshold for the downstream switch. C) Representative TNF-α production when providing costimulation to CD28 by recombinant biotinylated CD86 and averaged E_max_ values with SD from 4 independent experiments. D) Representative TNF-α production when providing costimulation to 4-1BB by recombinant biotinylated trimeric 4-1BBL and averaged E_max_ values with SD from 3 independent experiments. Additional cytokines are shown in Fig. S7-8. Statistical significance was determined by ordinary one-way ANOVA corrected for multiple comparisons by Dunnett’s test.

Activation-induced cell death may also result in reduced cytokine production but this is unlikely to be the case here. First, we previously confirmed that cell death is minimal in this experimental system (less than 10% of T cells stain positive for Annexin V at 8 hours) (47) and second, adapted T cells can be fully reactivated with co-stimulation (see below; Fig. 4C-D, transfer to pMHC + CD86 or 4-1BBL).

Taken together, perfect adaptation in cytokine production is observed with similar temporal kinetics across a 2,000-fold variation in antigen concentration and a 100,000-fold variation in antigen affinity.

### Perfect adaptation cannot be explained by complete TCR or pMHC downregulation

Previously, it has been shown that complete receptor downregulation can produce perfect adaptation provided that the receptor is not replenished (re-expressed) on the surface on the adaptation timescale (Fig. 1A). We therefore examined the surface dynamics of the TCR in our experimental system using flow cytometry. Consistent with previous reports (16, 18, 54–56), we observed concentration- and affinity-dependent TCR downregulation that reached steady-state within ∼3-6 hours (Fig. 2D). In all conditions tested, the TCR was only partially downregulated and this was not a result of a fraction of T cells downregulating their TCR (i.e. digital downregulation) because histograms showed the entire population of TCR-transduced T cells reducing their TCR surface expression (i.e. analogue downregulation, see Fig. 2D). Consistent with previous reports (18, 57), we observed a small but significant recovery in TCR surface expression on the timescale of ∼4 hours (Fig. 2E) suggesting that partial downregulation is maintained by a balance of re-expression and antigen-induced downregulation. Taken together, perfect adaptation cannot be explained by complete downregulation of the TCR.

It has also been shown that complete removal of the ligand can produce perfect adaptation (Fig. 1B). As already discussed, the efficient removal of all pMHC ligands is not known to take place during T cell activation with previous reports showing that pMHC ligands continually engage TCRs (16, 17). Indeed, the removal of pMHC is unlikely to be the mechanism in this experimental system because transferring T cells after they have adapted to plates newly coated with pMHC did not reactivate them to produce cytokine (see below; Fig. 4C-D, transfer to pMHC). Taken together, perfect adaptation by T cells cannot be explained by complete TCR or pMHC downregulation.

### Perfect adaptation by imperfect adaptation at the TCR coupled to a downstream switch

Given that up-regulation of the TCR can be observed on the adaptation timescale suggests that TCR downregulation would lead to imperfect adaptation (Fig. 1C). Therefore, additional mechanism(s) are required to produce perfect adaptation in cytokine production. Given that switches have been extensively documented in the TCR signalling pathways (17, 58–60) and that digital cytokine production has been reported (17), we hypothesised that a downstream switch could convert imperfect adaptation at the TCR into perfect adaptation in cytokine production (Fig. 1C).

To test this hypothesis, we converted the schematic (Fig. 1C) into an ordinary-differential-equation (ODE) model and used Approximate Bayesian Computations coupled to Sequential Monte Carlo (ABC-SMC) (61) to directly fit the model to the surface TCR and cytokine data (see Materials & Methods). Given that the experimental data is derived from a heterogeneous population of T cells, the ABC-SMC method is particularly appropriate because it effectively simulates a population of T cells with potentially different values of the model parameters representing population heterogeneity (Fig. S4).

The model produced an excellent fit to the data (Fig. 3A, solid lines) indicating that TCR downregulation coupled to digital cytokine production is sufficient to explain perfect adaptation. Importantly, the model reproduced perfect adaptation with partial TCR downregulation. By examining the timecourse at a single concentration (Fig. 3B), it was observed that incomplete TCR downregulation (R, surface TCR) lead to imperfect adaptation in TCR/pMHC complexes (C, receptor output). Perfect adaptation in cytokine production (O, output) was observed in the model because the concentration of TCR/pMHC complexes decreased below the switch threshold required to maintain the output.

**Figure 3:**
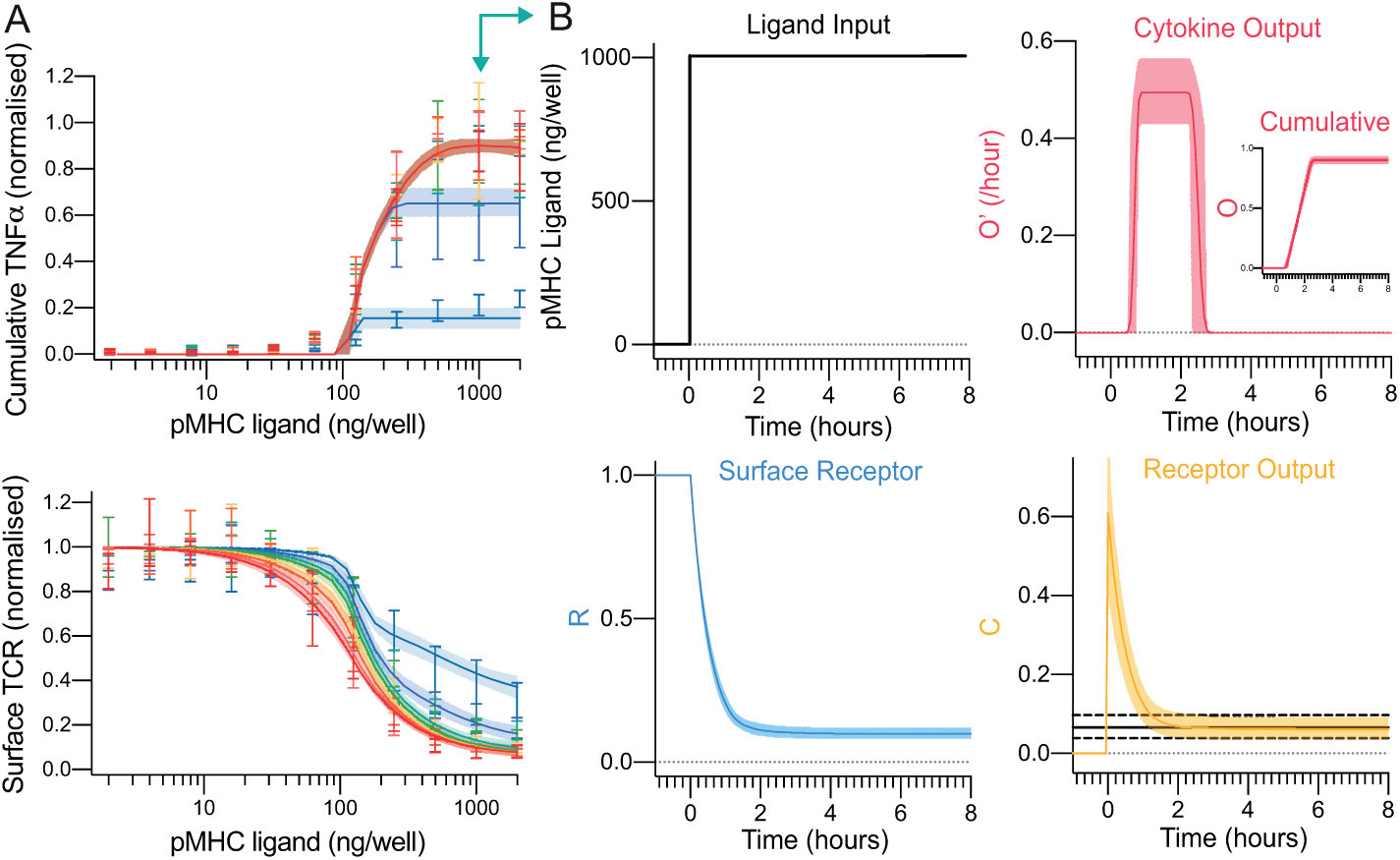
A mechanistic mathematical model shows that TCR downregulation coupled to a downstream switch is sufficient to explain perfect adaptation in T cell cytokine production to constant pMHC ligand stimulation. A) The fit of the mathematical model (Fig. 3C) using ABC-SMC to the physiological affinity pMHC data (Fig. 2B,D) with solid line and shaded region indicating the mean and 95% CI of the fit. B) Model outputs over time for a single concentration (1000 ng/well, teal arrow in panel A). The solid and dashed horizontal black lines in receptor output (bottom right) indicate the fitted mean threshold and 95% confidence intervals, respectively, for the downstream switch. Distributions of all fitted parameters can be found in Fig. S4.

This mechanism predicts that increasing the antigen strength by increasing its concentration or affinity would induce further TCR downregulation and cytokine production. Therefore, the model was used to predict the outcome of increasing the antigen concentration (Fig. S5) or affinity (Fig. S6) and experiments confirmed that TCR levels tuned to the new antigen strength with further cytokine production. As expected, reducing the antigen strength by reducing antigen affinity did not lead to marked changes in TCR expression or further cytokine production (Fig. S6D).

In summary, and in contrast to adaptation by other receptors, perfect adaptation can be explained here by imperfect adaptation at the TCR by partial downregulation coupled to a switch in the pathway for cytokine production (Fig. 1C).

### T cell adaptation to constant pMHC antigen can be overridden by costimulation

The model predicted imperfect adaptation by TCR downregulation so that residual TCR output continued after cytokine production had stopped (Fig. 3B). Given that T cells can encounter antigen *in vivo* with costimulation through other surface receptors, and costimulation is thought to lower the signalling threshold for cytokine production (62–64), we determined whether costimulation can amplify residual TCR signalling to reactivate adapted T cells.

We used the mathematical model to predict the outcome of transferring T cells from a first stimulation to a second stimulation on the same antigen with or without costimulation (Fig. 4A). Note that in these transfer experiments, T cells experience the same concentration of antigen in the first and second stimulation. The effect of costimulation was simulated by lowering the threshold of the switch required for cytokine production and as expected, this allowed T cells to produce cytokine provided they also continued to receive constant antigen stimulation (Fig. 4B).

In order to test whether CD28 costimulation could override adaptation, we stimulated T cells with the physiological affinity pMHC (first stimulation) before transferring them to the same titration of pMHC with or without recombinant CD86, which is the ligand for CD28 (second stimulation). Consistent with the adaptation phenotype, there was a dramatic reduction in TNF-α production in the second stimulation without CD86 but when CD86 was present, strong cytokine production was observed (Fig. 4C). Importantly, T cells transferred to empty wells without pMHC or to wells coated with only CD86 produced no cytokines.

In addition to CD28, the costimulatory receptor 4-1BB is also known to play an important role in the activation of CD8^+^ T cells. We repeated the experiments with the recombinant ligand to 4-1BB showing that this TNFR is also able to override adaptation but as with CD28, it critically relied on TCR/pMHC interactions (Fig. 4D).

Previous work on T cell anergy has described unresponsive T cell states that are induced when T cells are activated in the absence of CD28 costimulation. We therefore tested whether CD28 costimulation can prevent adaptation. We repeated the CD28 costimulation transfer experiments but now transferred T cells that were stimulated with either pMHC alone or with both pMHC and CD86 in the first stimulation to a second stimulation that included pMHC alone, CD86 alone, pMHC and CD86, or empty wells. We observed reduced cytokine production in the second stimulation to pMHC alone, which was similar to empty wells, irrespective of whether CD86 was included in the first stimulation (Fig. S8), suggesting that CD28 costimulation cannot prevent adaptation to constant antigen.

Taken together, these results indicate that perfect adaptation in cytokine production induced by constant pMHC antigen stimulation does not lead to perfect adaptation in TCR signalling because extrinsic costimulation through CD28 or 4-1BB can induce adapted T cells to produce TNF-α in a pMHC-dependent manner. This phenotype was also observed for other cytokines (Fig. S7-8).

### Adaptation by CAR-T cells to constant antigen can be overridden by costimulatory domains

Given that CAR-T cells experience constant antigen stimulation *in vivo*, we analysed their adaptation phenotype. To do this, we utilised the previously described T1 CAR (65) fused to the cytoplasmic tail of the ζ-chain (1^st^ generation CAR) that also recognises the NY-ESO-1_157−165_ peptide on HLA-A2 in a similar orientation to the TCR (K_D_= 4 nM (66)). These CAR-T cells were first stimulated with a titration of 9V pMHC before being transferred for a second stimulation on the same titration of 9V (Fig. 5A).

**Figure 5:**
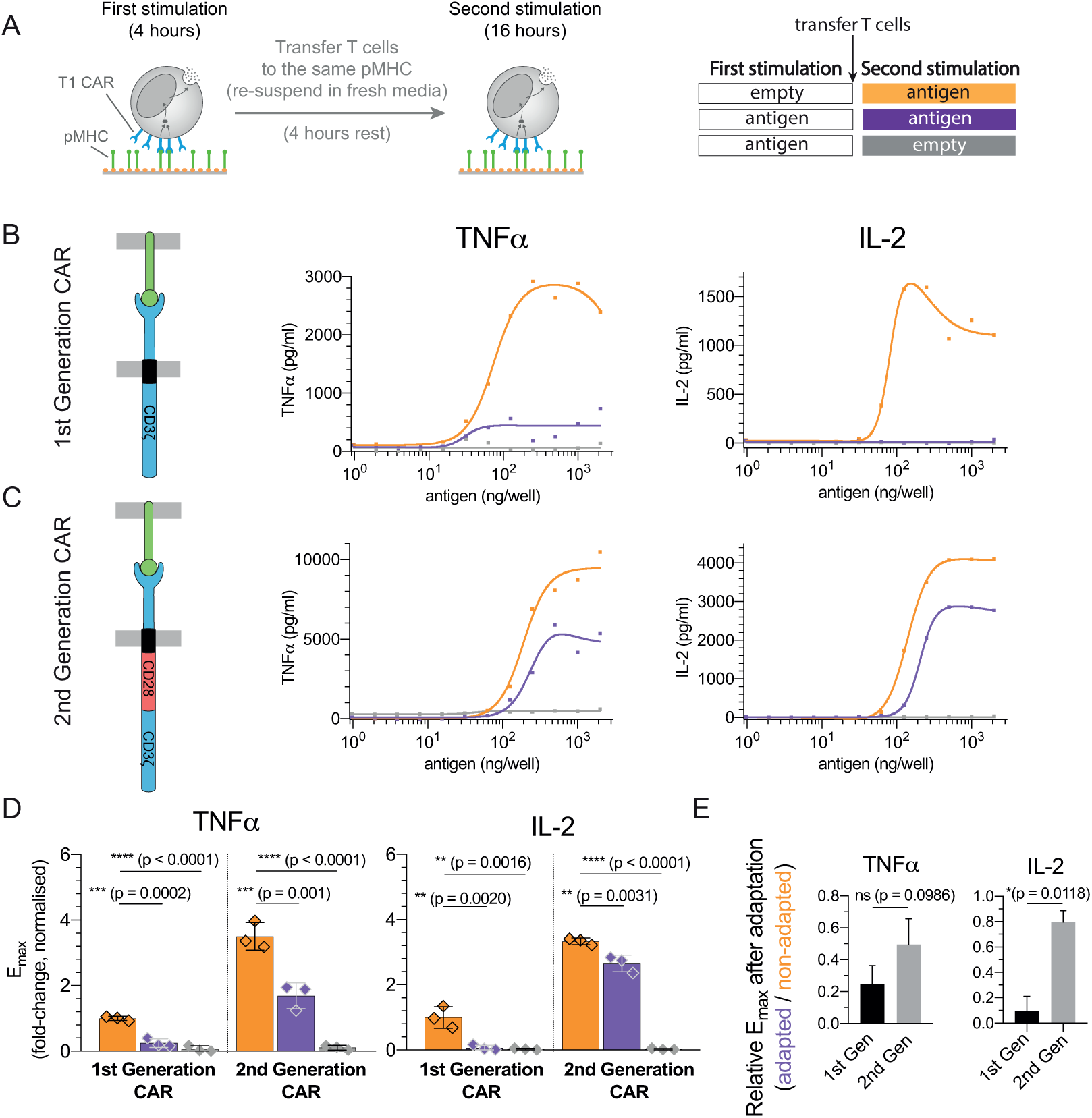
Adaptation is partially avoided in 2nd but not 1st generation CAR-T cells. A) Schematic of the experiment. T cells expressing the T1 CAR that recognises the 9V pMHC antigen were transferred to the same titration of antigen. B-C) Representative TNF-α and IL-2 production over antigen concentration from CAR-T cells expressing the B) the 1st generation variant containing only the ζ-chain and C) the 2nd generation variant containing the cytoplasmic tail of CD28 fused to the ζ-chain. D) Averaged E_max_ values and SD for 3 independent experiments. E) Fold reduction of E_max_ between the first and second stimulation for the 1st and 2nd generation CARs highlighting that 2nd generation CARs are more resistant to adaptation induced by constant antigen. Expression profile of both CARs and antigen-induced CAR downregulation is shown in Fig. S9. Statistical significance was determined by ordinary one-way ANOVA corrected for multiple comparisons by Dunnett’s test.

We observed reduced cytokine production by CAR-T cells that experienced the antigen in the first stimulation compared to CAR-T cells that were directly placed on the second stimulation (Fig. 5B, purple and orange lines, respectively). Given that costimulation can override adaptation, we repeated the experiments with a 2nd generation CAR containing the CD28 costimulatory domain finding that these CAR-T cells were able to partially avoid adaptation (Fig. 5C). Compared to cytokine production in the 1st stimulation (100%), the production of TNF-α and IL-2 were reduced in the second stimulation to 26% (p=0.0002) and 2.1% (p=0.002) in the 1st Generation CAR but were only reduced to 58.7% (p=0.001) and 79% (p=0.0031) in the 2nd generation CAR (Fig. 5D-E). We note that overall cytokine production was higher in the 2nd generation CAR (Fig. 5B-D) even though this receptor was consistently expressed at lower levels (Fig. S9).

Taken together, the adaptation phenotype observed with the TCR can also be observed with a 1st generation CAR that can be partially overridden by a 2nd generation CAR that includes costimulation. Partial rescue in the 2nd generation CAR is not unexpected because, unlike the complete rescue of the TCR by extrinsic co-stimulation (Fig. 4), co-stimulation in the CAR is intrinsic and is therefore reduced over time as a result of CAR downregulation.

## Discussion

Using a reductionist system to provide T cells with constant pMHC antigens, we observed that *in vitro* expanded primary human CD8^+^ T cells do not maintain cytokine production but instead exhibit perfect adaptation across a 100,000-fold variation in affinity. This adaptation can be rescued by increasing the antigen concentration (Fig. S5), affinity (Fig. S6), or when providing co-stimulation (Fig. 4).

### Mechanism of adaptation

Adaptation by surface receptors has been termed state-dependent inactivation (5, 9), which is an incoherent feedforward loop whereby ligand binding induces receptor signalling (positive regulation) and receptor downregulation (negative regulation) (7). Perfect adaptation takes place if the receptor is completely inactivated or downregulated by the ligand. Unlike other receptors, the TCR is only partially downregulated in response to antigen ligands leading to imperfect adaptation. To explain perfect adaptation in cytokine production, an additional downstream mechanism is required, and in the present study we have invoked a switch (Fig. 3C). However, other downstream mechanisms, such as additional IFFLs or NFLs, may also be able to convert imperfect adaptation at the TCR to perfect adaptation in cytokine production. Consistent with imperfect adaptation at the TCR, we found that extrinsic CD28 and 4-1BB costimulation can reactivate adapted T cells in a pMHC-dependent manner (Fig. 4).

Although the downstream switch could be replaced by an IFFL or NFL, models where these motifs are responsible for perfect adaptation and are downstream of the switch could not explain all adaptation rescue experiments. For example, a model where the sensitivity of the switch is sufficiently high so that it detected the presence of antigen independent of TCR levels could generate perfect adaptation if coupled to a downstream NFL or IFFL (Fig. S10). However, in this model, increasing the antigen strength cannot re-activate T cells after adaptation because the the input into the downstream NFL or IFFL remains unchanged (i.e. the switch remains in the on-state filtering out the analogue information on increasing antigen strength). Given that T cells can be reactivated by increasing the antigen strength (Fig. S6,S5), suggests that the analogue information of imperfect adaptation at the TCR is a central mechanism regulating T cell responses.

The minimal model of TCR downregulation coupled to a downstream switch can produce bell-shaped doseresponse curves (e.g. Fig. S6B). Previously, we argued that bell-shaped dose-response curves can be explained by incoherent feedforward loops but not by TCR downregulation (47). The key assumption underlying this conclusion was that the rate of cytokine production was in the steady-state, which is the case for Jurkat T cell lines (47), but the detailed analysis in the present work has revealed that it is not the case for primary T cells in the absence of costimulation. The bell-shaped dose-response curve produced by the kinetic model used here is a result of faster TCR downregulation at higher antigen concentrations resulting in cytokine production stopping at an earlier time point.

### Function of adaptation

Adaptation is a critically important and widely implemented process in biology. Unlike other receptors, perfect adaptation in T cells is achieved by imperfect adaptation at the TCR. This has the important consequence that adapted T cells are rendered dependent on both extrinsic costimulation and pMHC. In the specific example of activated T cells, whose killing capacity is thought to be less dependent on costimulation, perfect adaptation in cytokine production may be an important mechanism to ensure that their ability to initiate or maintain inflammation is extrinsically regulated by other cells. In this way, perfect adaptation may serve to balance functional immunity with excessive tissue damage.

### Relation to *in vivo* studies

T cells are known to enter unresponsive states upon recognition of persistent self- and viral-antigens *in vivo* (2, 3, 20, 31, 33–38, 67–71). While the underlying mechanisms that induce and maintain these states are debated, their functional phenotype is characterised by transient cytokine production that can be overcome by costimulation as observed here (Fig. 2,4). For example, it has been shown that effector CD4^+^ T cells transiently produce cytokines despite continued antigen exposure (70) and the unresponsive (exhausted) phenotype of CD8^+^ T cells induced by persistent antigen stimulation can be overcome by costimulation (71). We note that the mechanism of adaptation that we report can take place with only minor TCR downregulation, which may help reconcile previous reports arguing either that TCR downregulation can explain tolerance (20, 30–33) or that TCR downregulation did not play a role in tolerising T cells because overt downregulation was not observed (34–38). On the other hand, our results suggest that adaptation by TCR downregulation renders T cells dependent on costimulation, which is in line with the finding that T cells with impaired TCR downregulation lose their dependence on costimulation for activation (25, 27).

Ultimately, by studying *in vitro*-expanded human T cells, we are inherently limited in making direct comparisons with *in vivo* T cell phenotypes. As methods for the generation of large numbers of antigen specific human T cells improve, it would be important to examine the response of natural quiescent T cell populations (e.g. naive CD8 and CD4 T cells) in this experimental assay.

### Implications for adoptive therapy

The T cells we have used were generated using a protocol for adoptive therapy with TCRs or CARs. Consistent with the adaptation phenotype we observe, it has recently been observed that CAR-T cells exposed to chronic antigen become unresponsive but can respond to a higher antigen dose, which correlates with CAR expression (43). The ability of 2nd generation CARs to partially avoid adaptation in cytokine production (Fig. 5) may explain why they generate much more potent and persistent anti-tumour responses *in vivo* (72–75) even though their *in vitro* killing capacity is comparable to 1st generation receptors (73–76). The optimisation of TCRs and CARs has focused on affinity, surface levels, and signalling potency, but engineering for optimal downregulation has yet to be explored.

## Materials & Methods

### Protein production

pMHCs were refolded *in vitro* from the extracellular residues 1-287 of the HLA-A*02:01 α-chain, β2-microglobulin and NY-ESO-1_157−165_ peptide variants as described previously (47). CHO cell lines permanently expressing the extracellular part of human CD86 (amino acids 6-247) with a His-tag for purification and a BirA biotinylation site were kindly provided by Simon Davis (Oxford, UK). Cells were cultured in GS selection medium and passaged every 3-4 days. After 4-5 passages from thawing a new vial, cells from 2 confluent T175 flasks were transferred into a cell factory and incubated for 5-7 days after which the medium was replaced. The supernatant was harvested after another three weeks, sterile filtered and dialysed over night. The His-tagged CD86 was purified on a Nickel-NTA Agarose column. 4-1BB Ligand expression constructs were a kind gift from Harald Wajant (Wuerzburg, Germany) and contained a Flag-tag for purification and a tenascin-C trimerisation domain. We added a BirA biotinylation site. The protein was produced by transient transfection of HEK 293T cells with XtremeGene™ HP Transfection reagent (Roche) according to the manufacturer’s instructions and purified following a published protocol (77), with the exception of the elution step where we used acid elution with 0.1 M glycine-HCl at pH 3.5. The pMHC or costimulatory ligand was then biotinylated *in vitro* by BirA enzyme according to the manufacturer’s instructions (Avidity) and purified using size-exclusion chromatography.

### Production of lentivirus for transduction

HEK 293T cells were seeded into 6-well plates 24 h before trans-fection to achieve 50–80% confluency on the day of transfection. Cells were cotransfected with the respective third-generation lentiviral transfer vectors and packaging plasmids using Roche XtremeGene™ 9 (0.8µg lentiviral expression plasmid, 0.95 µg pRSV-rev, 0.37 µg pVSV-G, 0.95 µg pGAG). The supernatant was harvested and filtered through a 0.45 µm cellulose acetate filter 24-36h later. The 1G4 TCR used for this project was initially isolated from a melanoma patient (78). The affinity maturation to the c58c61 TCR variant used herein was carried out by Adaptimmune Ltd. The TCR and all CARs in this study have been used in a standard third-generation lentiviral vector with the EF1α promoter. The CAR constructs that bind the NY-ESO-1_157−165_ HLA-A2 pMHC complex (66, 79) were a kind gift from Christoph Renner (Zurich, Switzerland). The high-affinity T1 version was used for this project. All CAR constructs contained the scFv binding domain, a 2 Ig domain spacer derived from an IgG antibody Fc part and the CD28 transmembrane domain. We modified the different CARs to contain the CD3ζ signalling domain alone or in combination with the CD28 signalling domain.

### T cell isolation and culture

Up to 50 ml peripheral blood were collected by a trained phlebotomist from healthy volunteer donors after informed consent had been taken. This project has been approved by the Medical Sciences Inter-Divisional Research Ethics Committee of the University of Oxford (R51997/RE001) and all samples were anonymised in compliance with the Data Protection Act. Alternatively, leukocyte cones were purchased from National Health Services Blood and Transplant service. Only HLA-A2^-^ peripheral blood or leukocyte cones were used due to the cross-reactivity of the high-affinity receptors used in this project which leads to fratricide of HLA-A2^+^ T cells (65, 66, 80). CD8^+^ T cells were isolated directly from blood using the CD8^+^ T Cell Enrichment Cocktail (StemCell Technologies) and density gradient centrifugation according to the manufacturer’s instructions. The isolated CD8^+^ T cells were washed and resuspended at a concentration of 1 × 10^6^ cells per ml in completely recon-stituted RPMI supplemented with 50 units/ml IL-2 and 1 × 10^6^ CD3/CD28-coated Human T-Activator Dynabeads (Life Technologies) per ml. The next day, 1 × 10^6^ T cells were transduced with the 2.5 ml virus-containing supernatant from one well of HEK cells supplemented with 50 units/ml of IL-2. The medium was replaced with fresh medium containing 50 units/ml IL-2 every 2–3 d. CD3/CD28-coated beads were removed on day 5 after lentiviral transduction and the cells were used for experiments on days 10-14. This protocol produces antigen-experienced CD8^+^ T cells with a fraction (typically ∼70%) expressing the transduced c58c61 TCR or T1 CAR.

### T cell stimulation

T cells were stimulated with titrations of plate-immobilised pMHC ligands with or without co-immobilised ligands for accessory receptors. Ligands were diluted to the working concentrations in sterile PBS. 50 µl serially two-fold diluted pMHC were added to each well of Streptavidin-coated 96-well plates (15500, Thermo Fisher). After a minimum 45 min incubation, the plates were washed once with sterile PBS. Where accessory receptor ligands were used, those were similarly diluted and added to the plate for a second incubation of 45-90 min. In experiments with small molecule inhibitors, the T cells were incubated with the inhibitor at 37 °C for 20-30 min prior to the start of the stimulation. The inhibitors were left in the medium for the whole duration of the stimulation. All control conditions were incubated with DMSO at a 1:1000 dilution so that the DMSO concentration was the same for inhibitor and non-inhibitor samples. PP2 was used at a 20 µM concentration. After washing the stimulation plate with PBS, 7.5 × 10^4^ T cells were added in 200 µl complete RPMI without IL-2 to each stimulation condition. The plates were spun at 9 x g for 2 min to settle down the cells and then incubated at 37 °C with 5 % CO_2_. For transfer experiments, the T cells were pipetted from the stimulation plate into a V-bottom plate and pelleted after the first round of stimulation. The supernatant was stored at −20 °C for later cytokine ELISAs, the cells were resuspended in 200 µl fresh R10 medium and – depending on the experiment – either rested for some time or transferred to another stimulation plate with a new set of conditions. The cells were then again settled down by centrifuging at 9 x g before incubation.

### Flow cytometry

Flow cytometry was used to assess receptor expression after TCR and CAR transductions, and to quantify receptor downregulation at the end of stimulation experiments. After stimulation, T cells were pelleted in a V-bottom plate and resuspended in 40 µl PBS with 2% BSA and fluorescent 9V pMHC tetramers that were produced with Streptavidin-PE (Biolegend, 405204) and used at a predetermined dilution (1:100-1:1000). The staining was incubated for 20-60 min after which the cells were pelleted, resuspended in 70-100 µl PBS and analysed on a FACSCalibur™ or LSRFortessa X-20 (BD) flow cytometer. Flow cytometry data was analysed with Flowjo V10.0.

### ELISA

Supernatants from stimulation experiments were stored at −20 °C. Cytokine concentrations were measured by ELISAs according to the manufacturer’s instructions in Nunc MaxiSorp™ flat-bottom plates (Invitrogen) using Uncoated ELISA Kits (Invitrogen) for TNF-α, IFN-γ, MIP-1β, and IL-2.

### Data analysis

The fraction of T cells expressing the transduced TCR or CAR and the amount of supernatant cytokine exhibited variation between independent experiments (with different donors). Therefore, directly averaging the data produced curves that were not representative of individual repeats. To average cytokine data, the maximum for each repeat was normalised to 1 before averaging independent repeats. To average surface TCR gMFI (*X*), which is on a logarithmic scale, we used the following formula that corrected for the fraction of T cells expressing the TCR 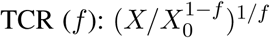, where *X*_0_ is the mean background gMFI of the TCR negative population. After applying this transform, each repeat was normalised to the maximum gMFI before averaging independent repeats.

The statistical analysis of maximum cytokine produced across different pMHC concentrations (E_max_), was performed by expressing *E*_max_ as a fold-change to pMHC alone before averaging independent repeats. Given that the dose-response curves often exhibited a bell-shape, it was not possible to use a standard Hill function to estimate E_max_. Instead, we used *lsqcurvefit* in Matlab (Mathworks, MA) to fit a function that was the difference of two Hill curves in order to generate a smooth spline through the data from which the maximum value of cytokine was estimated. This procedure was used to extract E_max_ values in Fig. 4, 5, S7, S8. In a limited number of cases, individual outlier values were excluded prior to data fitting but are still shown as data points in respective figures.

### Statistical analysis

Ordinary one-way ANOVA corrected for multiple comparisons by Dunnett’s test was performed on experimental data to determine statistical significance levels (Fig. 4C,D, Fig. S6C, Fig. 5D,E, Fig. S8). Statistical significance for surface TCR recovery (Fig. 2E) was performed by using an F-test for the null hypothesis that a single Hill curve (with the same parameters) can explain the data. GraphPad Prism was used for all statistical analyses.

### Mathematical model

The mathematical model (Fig. 1C) is represented as a system of two non-linear coupled ordinary-differential-equations (ODEs),

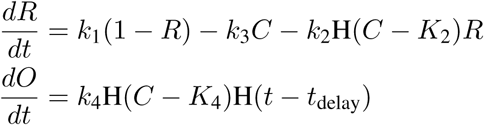

where *R* and *O* are the surface TCR levels (normalised to 1) and cumulative cytokine output with initial conditions 1 and 0, respectively. Given that TCR/pMHC binding kinetics (seconds) are faster than experimental timescales (hours), the concentration of TCR/pMHC complexes (*C*, defined as receptor output) were assumed to be in quasi steady-state, *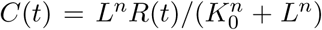*, where *L* is the given concentration of pMHC (in ng/well), *K*_0_ is the effective dissociation constant (in ng/well), and *n* is the Hill number. In the equation for *R*, the first two terms are the basal turnover of surface TCR (*k*_1_(1 − *R*)) and the pMHC binding induced downregulation of TCR (−*k*_3_*C*). In the equation for *O*, the term for the switch (*k*_4_H(*C* − *K*_4_)) includes a heaviside step function (H), so that the term is 0 unless receptor output (*C*) is above the switch threshold (*K*_4_), and in this case, cytokine output is produced at rate *k*_4_.

To directly fit the model to the data, two additional modifications were required. First, TCR downregulation is biphasic in time (55, 57) (e.g. Fig. S5) requiring an additional term for downregulation in the equation for *R* (*k*_2_H(*C* − *K*_2_)*R*). This term increases TCR downregulation initially when the output from the receptor (*C*) is above a threshold (*K*_2_) and at a molecular level, this may represent signalling-dependent bystander downregulation Given that the model already contained a stif step-function in the equation for *O*, this step-function was approximated by a Hill number with large cooperativitiy for computational efficiency 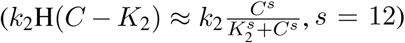. Second, cytokine production was larger in the 2nd hour compared to the 1st hour (e.g. Fig. 2B), which may be associated with transcriptional delays. To capture this delay, a multiplicative term in the equation for *O* was introduced (H(*t* − *t*_delay_)) so that cytokine production was only initiated after a delay of *t*_delay_.

### Data fitting using ABC-SMC

A Matlab (Mathworks, MA) implementation of a previously published algorithm for Approximate Bayesian Computation coupled to Sequential Monte Carlo (ABC-SMC) was used for data fitting (61). The ODEs were evaluated using the Matlab function *ode23s*.

Using ABC-SMC, the values of *R*(*t*) and *O*(*t*) were directly fitted to the normalised surface TCR levels and cumulative cytokine output, respectively, for the dose-response timecourse of 4A8K (192 data points in total). The distance measure was the standard sum-squared-residuals (SSR) and all 9 model parameters were fitted (*K*_0_, *n, k*_1_, *k*_2_, *K*_2_, *k*_3_, *k*_4_, *K*_4_, *t*_delay_). A population of 3000 particles were initialised with uniform priors in log-space and propagated through 30 populations by which point the distance measure was no longer decreasing. The final population of 3000 particles, each of which had a different set of model parameters (Fig. S4), was used to display the quality of the fit (Fig. 3A). Although the ODE model represents the reactions within a single cell, and hence the dose-response curve for a single cell would exhibit a perfect switch (i.e. a step function), the population averaged dose-response curves from the model include particles (i.e. cells) with different parameter values, accounting for population heterogeneity, leading to a more gradual dose-response curve.

The posterior distributions revealed that only a subset of the model parameters were uniquely determined (Fig. S4). Nonetheless, we were still able to make predictions using the model by simulating different experimental conditions for the 3000 particles in the final population. To predict the effect of increasing antigen concentration (Fig. S5B), the concentration of antigen was increased at the indicated times to the indicated value (no additional parameters were required). To predict the effect of increasing antigen affinity (Fig. S6B), the TCR/pMHC binding parameters in the model (*K*_0_ and *n*) for each particle were reduced by 50%. To predict the effect of co-stimulation (Fig. 4B), the threshold for the switch (*K*_2_) for each particle was reduced by 60% for a duration of 8 hours. The value of 60% was chosen as it approximately reproduced EC_50_ ∼ 100 ng/well observed in the data. The duration of this reduction scaled the value of E_max_ with 8 hours producing a value more similar to the 4-1BB and 16 hours producing a value more similar to the CD28 (not shown).

## Acknowledgements

We thank Simon J. Davis for providing CD86 expression plasmids, Harald Wajant for providing 4-1BBL expression plasmids, Christopher Renner for providing T1 CAR plasmids, Alan Rendall, Vahid Shahrezaei, and Eduardo Sontag for feedback on mathematical modelling, Adaptimmune Ltd for providing the c58c61 TCR, and Enas Abu Shah, Michael L. Dustin, Marion H. Brown, and Vincenzo Cerundolo for helpful discussions about experimental protocols. We thank P. Anton van der Merwe for a critical reading of the manuscript.

## Author contributions

NT, PK, SG, JN, and JP performed experiments; NT, PK, and OD performed the mathematical modelling; NT, PK, SG, JN, JP, and OD analysed data; NT, PK, and OD designed the research and wrote the paper; NT, PK, and SG contributed equally. All authors discussed the results and commented on the paper.

## Funding

This work was supported by a Doctoral Training Centre Systems Biology studentship from the Engineering and Physical Sciences Research Council (to NT), a scholarship from the Konrad Adenauer Stiftung (to NT), a studentship from the Edward Penley Abraham Trust and Exeter College, Oxford (to PK), a postdoctoral extension award from the Cellular Immunology Unit Trust (to PK), a Sir Henry Dale Fellowship jointly funded by the Wellcome Trust and the Royal Society (098363, to OD), pump-prime funding from the Cancer Research UK Oxford Centre Development Fund (CRUKDF 0715, to OD), National Science Foundation (USA) grant (1817936, to EDS) and a Wellcome Trust Senior Research Fellowship (207537/Z/17/Z, to OD).

## Supplementary Information

**Figure S1:**
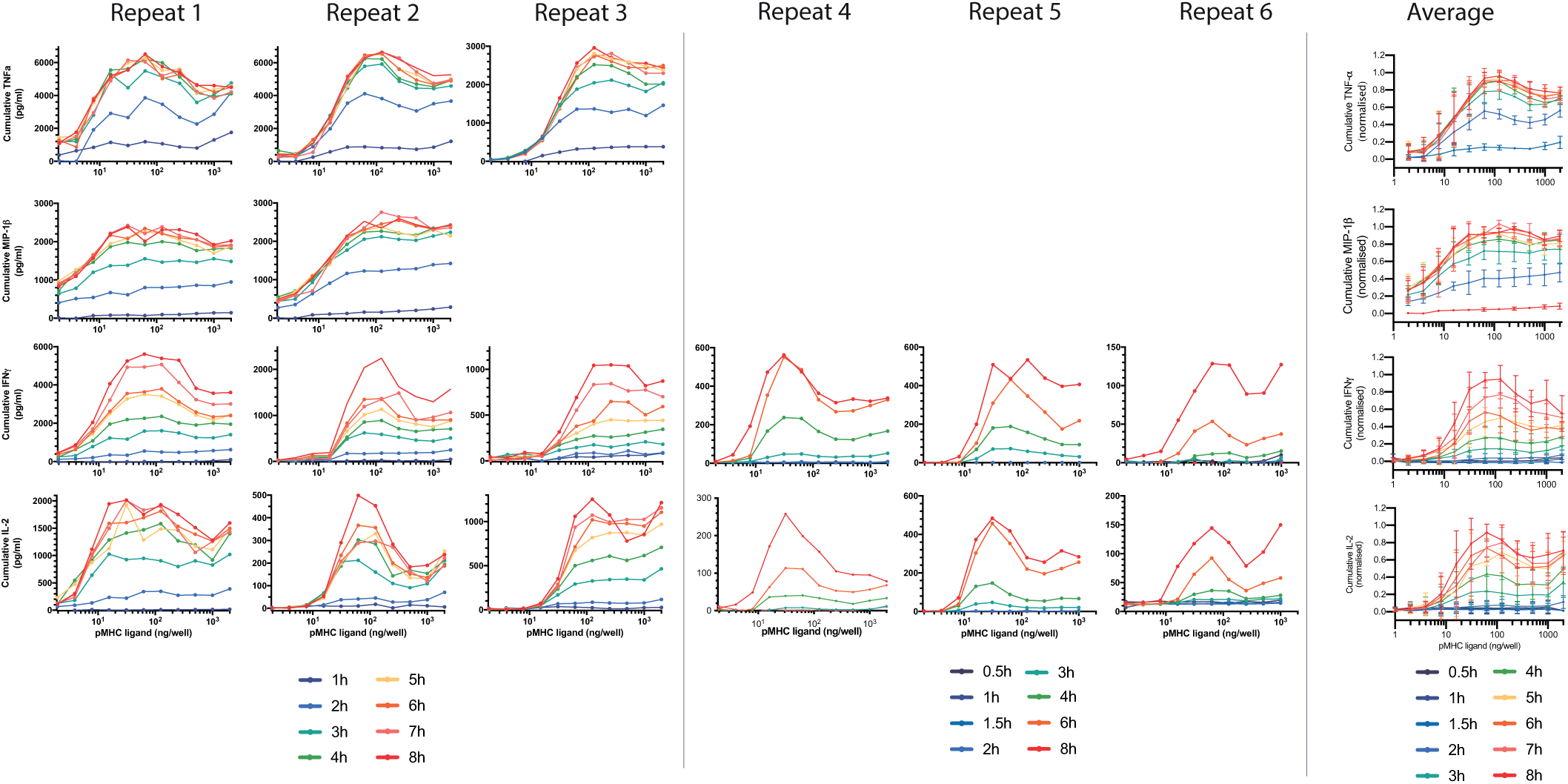
Cytokine production by T cells in response to constant stimulation by the supra-physiological affinity 9V pMHC. Expanded data from Fig. 2B showing TNF-α, MIP-1β, IFN-γ and IL-2 for individual repeats (6 left columns) along with averaged data (right column). The averaged TNF-α data is the same as in Fig. 2B. Error bars represent the SD.

**Figure S2:**
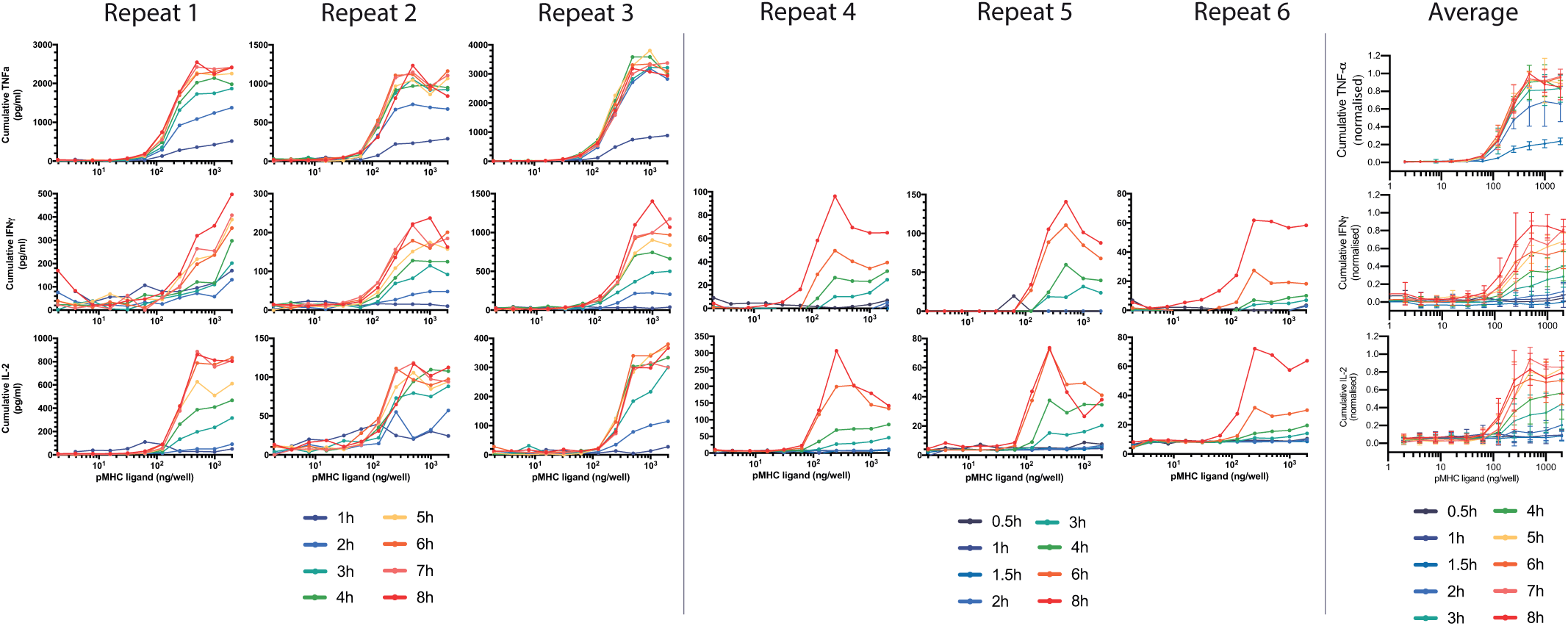
Cytokine production by T cells in response to constant stimulation by the physiological affinity 4A8K pMHC. Expanded data from Fig. 2B showing TNF-α, IFN-γ and IL-2 for individual repeats (6 left columns) along with averaged data (right column). The averaged TNF-α data is the same as in Fig. 2B. Error bars represent the SD.

**Figure S3:**
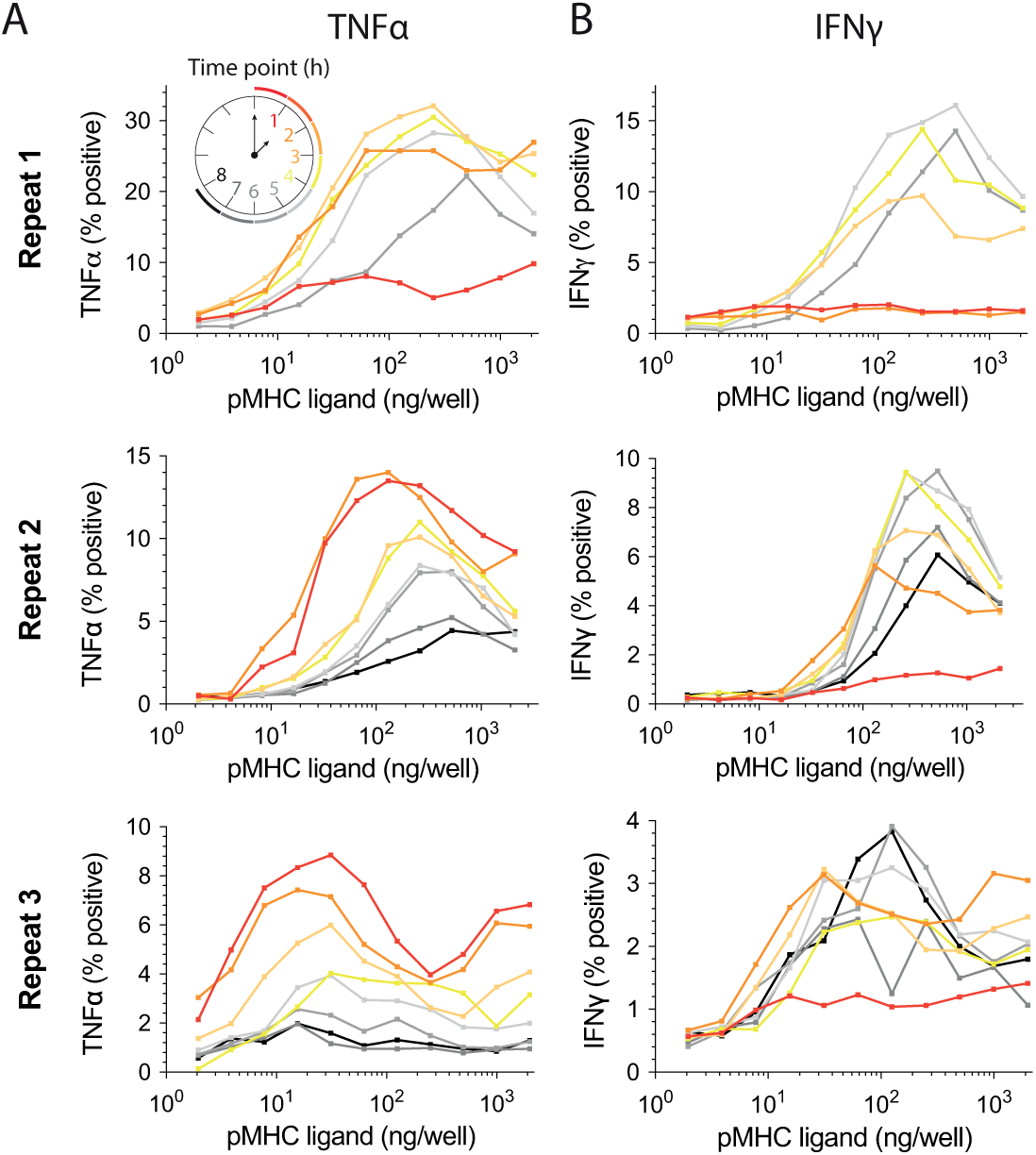
T cell adaptation is the result of reduced per-cell cytokine production. T cells were stimulated with the indicated concentration of 9V pMHC for the indicated time (1–8 hours) with intracellular cytokine staining of A) TNF-α and B) IFN-γ performed using flow cytometry. Cytokine secretion was blocked by addition of Brefeldin A for the last hour of the stimulation.

**Figure S4:**
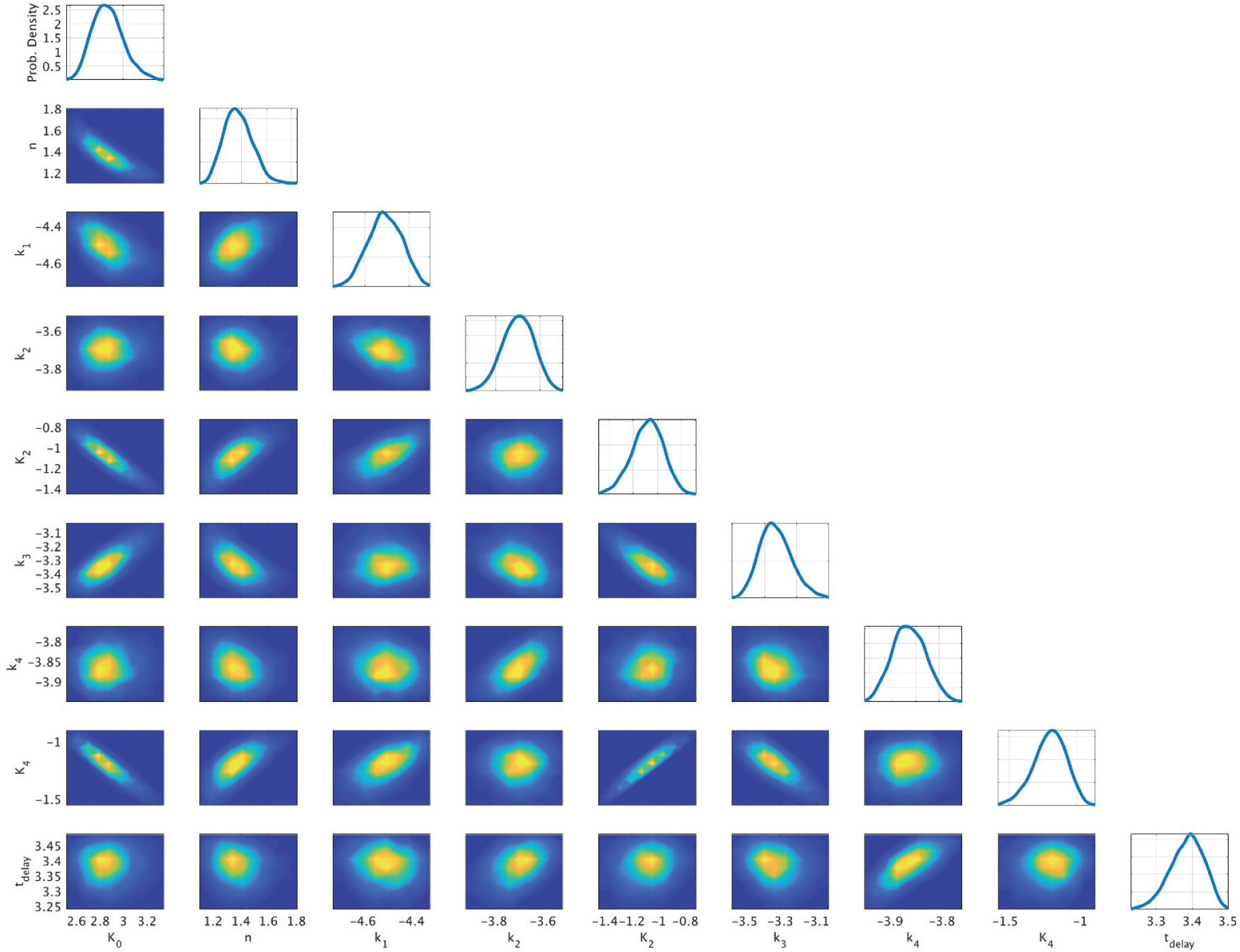
Posterior distributions of 3000 particles after 30 populations using ABC-SMC (see Materials & Methods). The marginal distribution of each parameter is shown on the diagonal with off-diagonal heatmaps displaying pair correlations.

**Figure S5:**
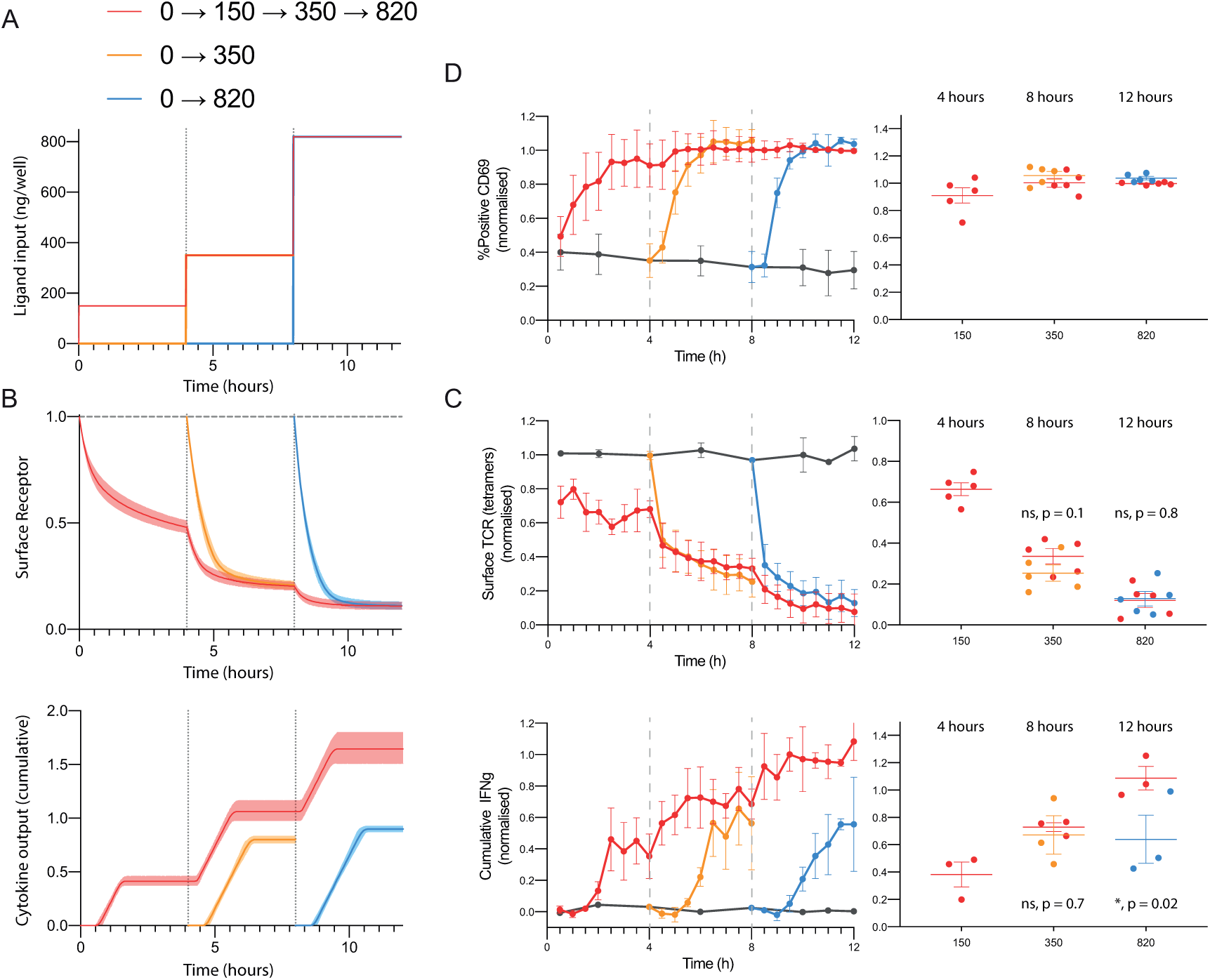
Increasing antigen concentration tunes TCR levels and induces further cytokine production. A) Schematic depicting the concentration of antigen (y-axis) experienced by three population of T cells (red, orange, blue) over time (x-axis). In the red population, T cells are initially placed on 150 ng/well of pMHC before being transferred at 4 hours to 350 ng/well and then at 8 hours to 820 ng/well (2.3-fold increases in concentration). In the case of the orange and blue populations, T cells were placed directly on 350 ng/well and 820 ng/well at 4 and 8 hours respectively. B) Predicted TCR surface expression (top) and cumulative cytokine production (bottom) by the mathematical model (see Materials & Method) highlights that increasing the antigen concentration leads to the tuning of TCR levels to the new antigen concentration with further cytokine production. C) Experimentally measured TCR surface levels (top) and cumulative IFN-γ (bottom) with statistical comparisons (right) confirms the model predictions. Statistics are performed by a paired t-test with p-values as indicated. D) Percent of T cells positive for CD69%. Data is shown as mean and SD from *>*4 independent experiments.

**Figure S6:**
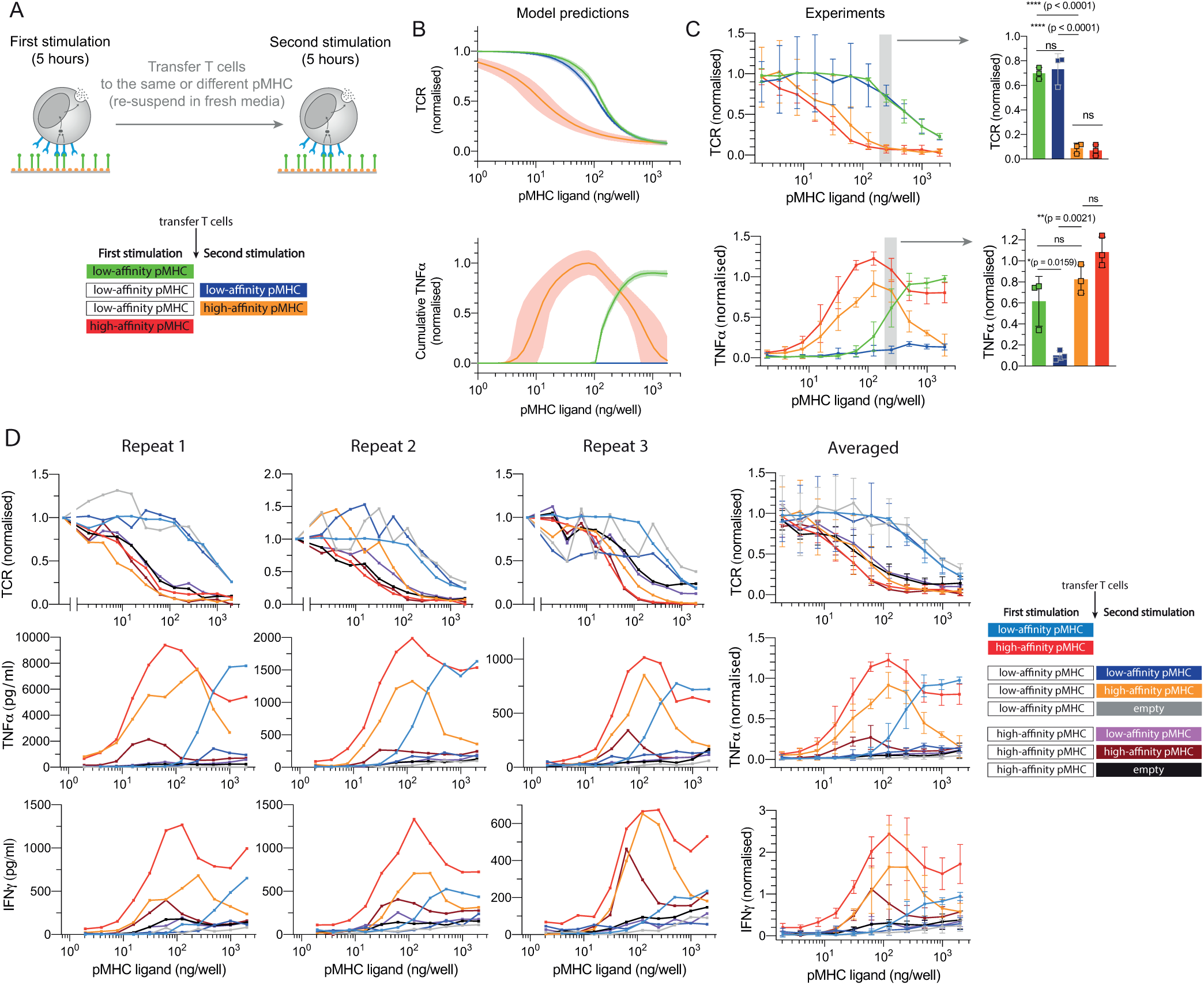
T cells adapted to a low-affinity antigen can be reactivated with a higher-affinity antigen. A) Schematic of the experiment showing that T cells were first stimulated for 5 hours on the low-affinity antigen before being transferred for a second stimulation of 5 hours with the same or different pMHC antigen (at the same antigen concentration). B) Predicted TCR surface expression (top) and cytokine production (bottom) for the transfer experiment by the mathematical model. C) TCR surface expression (top) and TNF-α production (bottom) with a detailed comparison performed at the indicated concentration (right). Consistent with the adaptation phe-notype, a first stimulation with the low-affinity 4A8K pMHC (green) leads to reduced cytokine production in a second stimulation on the same pMHC (blue). However, transferring T cells to the higher-affinity 9V pMHC leads to further TCR downregulation and further cytokine production (orange). It is noteworthy that beyond the grey shaded region cytokine production is progressively reduced, which the model predicts is a result of lower levels of surface TCR prior to transfer to the higher affinity pMHC. Data are mean and standard deviation of 3 independent experiments with statistical significance determined by ordinary one-way ANOVA corrected for multiple comparisons by Dunnett’s test. D) Individual repeats (left 3 columns) and averaged data (right column) showing TCR, TNF-α, and IFN-γ for all transfer conditions (see legend on right), including transfers from the high-affinity (9V) to the low-affinity (4A8K) pMHC showing that decreasing antigen affinity does not induce cytokine production (light purple). The averaged data in panel C is taken from the averaged data in panel D and shows a subset of all conditions tested.

**Figure S7:**
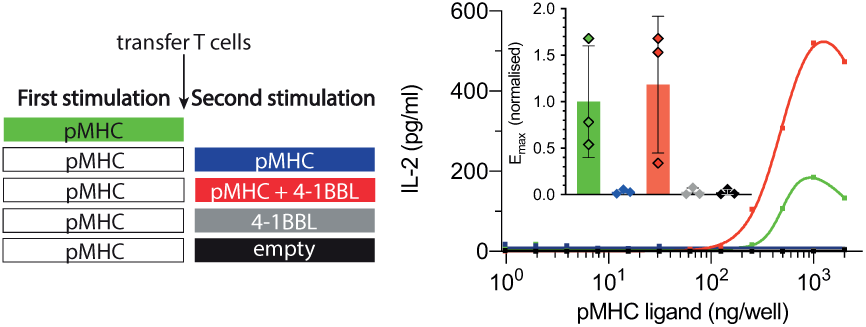
T cells adapted to constant antigen can be reactivated with 4-1BB costimulation. Expanded data showing IL-2 for the experiment in Fig. 4D.

**Figure S8:**
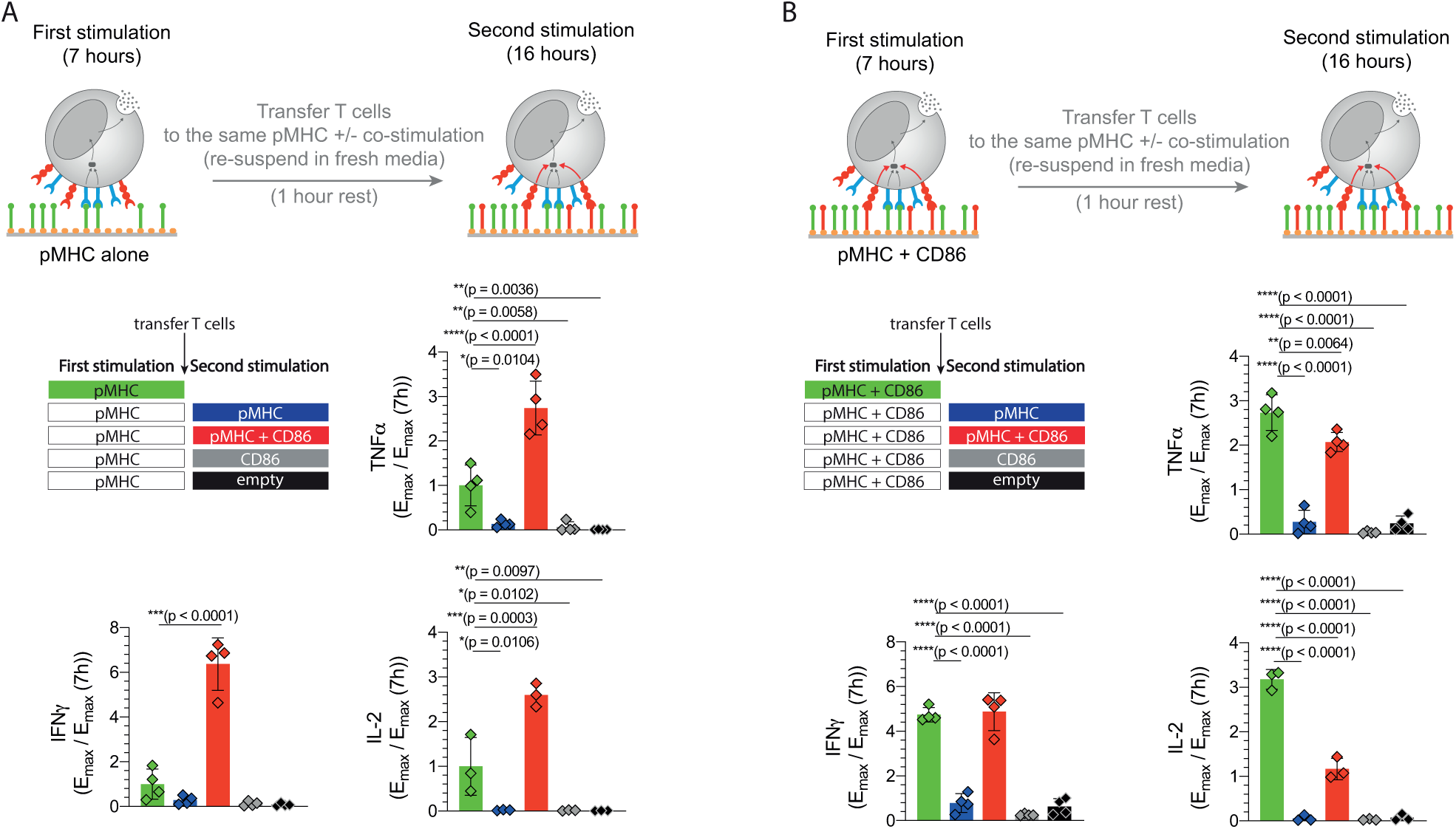
CD28 costimulation does not prevent adaptation to constant antigen stimulation. Primary human CD8^+^ effector T cells expressing the c58c61 TCR were stimulated with a concentration titration of the low-affinity pMHC antigen 4A8K and the supernatant concentration of TNF-α, IFN-γ and IL-2 measured by ELISA. T cells were first stimulated for 7h with the 4A8K low-affinity pMHC with or without recombinant CD86 (light blue lines). The T cells were rested for 1 hour in fresh medium before being transferred in fresh medium to pMHC alone (dark blue), pMHC and CD86 (red), CD86 alone (grey), or to empty wells (black). A) Schematic of transfer experiment with pMHC alone in the first stimulation (top) along with averaged E_max_ values from four independent experiments for TNF-α, IFN-γ, and IL-2 (bottom). B) Schematic of transfer experiment with pMHC and CD86 in the first stimulation (top) along with averaged E_max_ values from four independent experiments for TNF-α, IFN-γ, and IL-2 (bottom). All data is normalised to the value of E_max_ at 7 hours with pMHC alone. Error bars represent the SD and statistical significance was determined by ordinary one-way ANOVA corrected for multiple comparisons by Dunnett’s test comparing all conditions to the value of *E*_max_ during the first stimulation.

**Figure S9:**
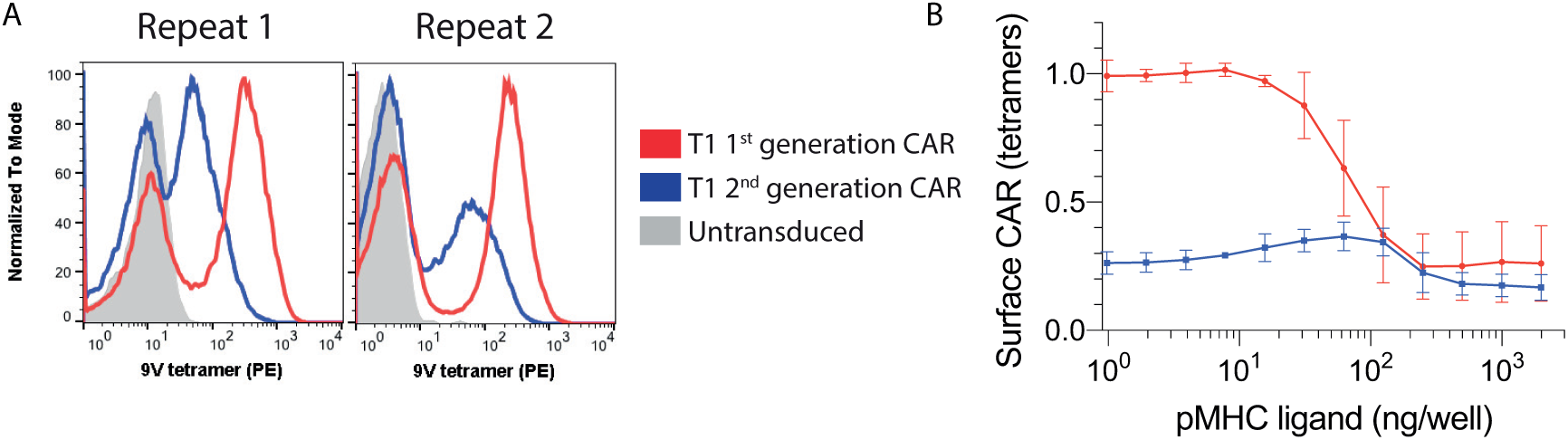
Expression levels and antigen-induced downregulation of 1st and 2nd generation CARs. A) T cells expressing the 1st and 2nd generation T1 CAR were stained with 9V pMHC tetramers and CAR expression was measured by flow cytometry. B) CAR-T cells were stimulated with the indicated titration of the 9V pMHC antigen for 8 hours with surface CAR expression measured using pMHC tetramers. Data is average of at least 3 independent experiments and error bars represent SD.

**Figure S10:**
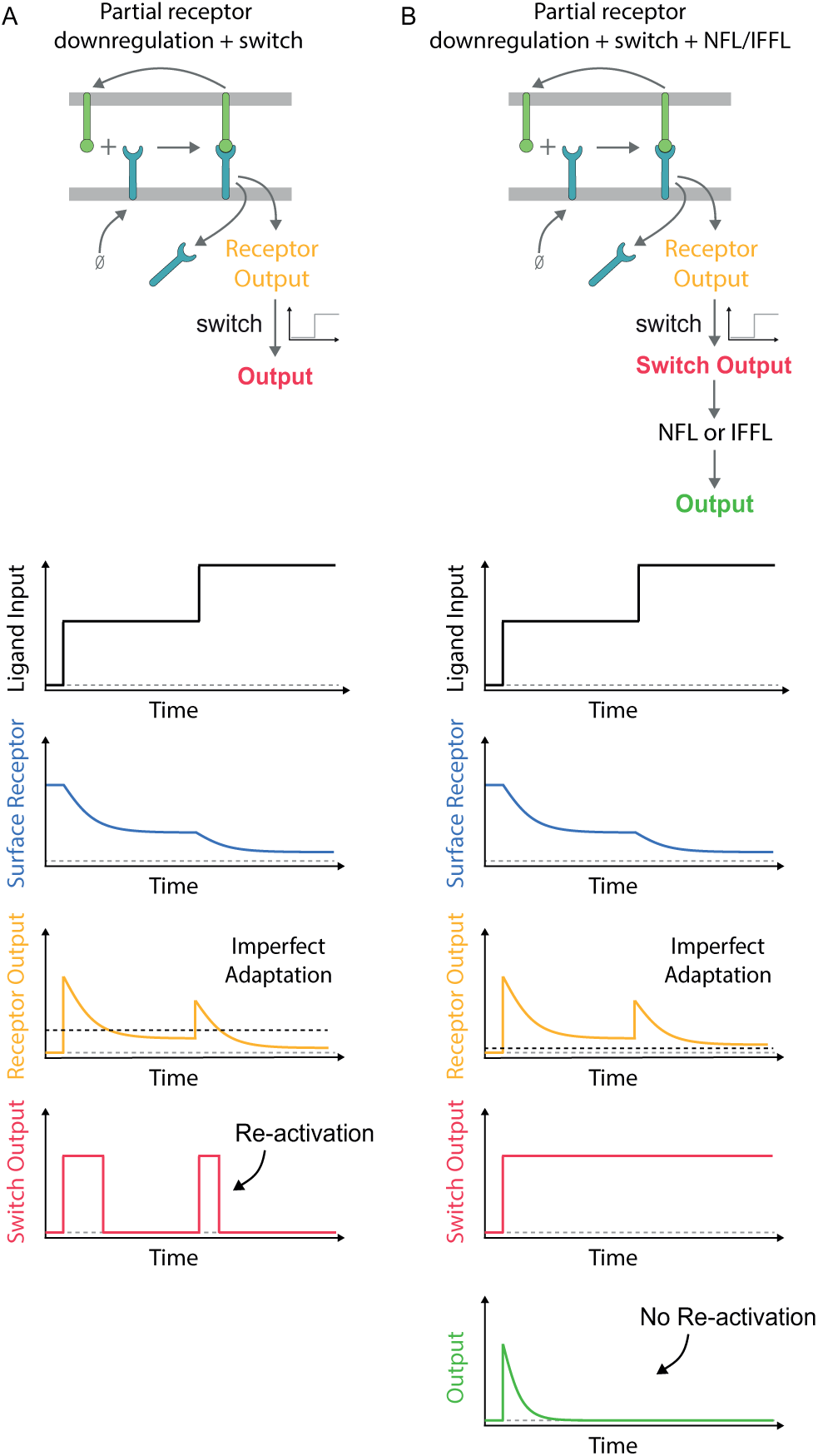
A modified model where TCR surface levels do not contribute to shaping cytokine production, as a result of a highly sensitive switch, cannot explain re-activation of T cells to higher antigen strength. A) As discussed in the main text and shown experimentally (Fig. S6, S5), the proposed model of imperfect adaptation at the TCR by downregulation coupled to a downstream switch predicts that increasing antigen strength can reactivate adapted T cells. B) On the other hand, increasing the sensitivity of the switch (i.e. lowering the threshold, horizontal dashed line on receptor output) so that it remains in the on-state whenever antigen ligand is present makes TCR surface levels, and hence adaptation, have no impact on downstream signalling. Perfect adaptation in this model now requires a downstream negative feedback loop (NFL) or incoherent feedforward loop (IFFL). However, in this model, re-activation of T cells by increasing the ligand strength is not possible because the analogue information on ligand strength is filtered out by the sensitive switch.

## Notes

#### Summary of Updates

Key changes: 1. Measurements of TCR downregulation were repeated with a more sensitive detection reagent resolving changes in TCR surface levels on the timescale of hours. 2. Supplementary data showing the effect of Lck inhibition on surface TCR levels were removed. 3. The mathematical model was simplified as a result of changes 1 and 2 and re-fit to all data. 4. Experiments showing that increasing antigen concentration can increase cytokine production are added.

